# A sister lineage of the *Mycobacterium tuberculosis complex* discovered in the African Great Lakes region

**DOI:** 10.1101/2020.01.20.912998

**Authors:** Jean Claude Semuto Ngabonziza, Chloé Loiseau, Michael Marceau, Agathe Jouet, Fabrizio Menardo, Oren Tzfadia, Rudy Antoine, Esdras Belamo Niyigena, Wim Mulders, Kristina Fissette, Maren Diels, Cyril Gaudin, Stéphanie Duthoy, Willy Ssengooba, Emmanuel André, Michel K Kaswa, Yves Mucyo Habimana, Daniela Brites, Dissou Affolabi, Jean Baptiste Mazarati, Bouke Catherine de Jong, Leen Rigouts, Sebastien Gagneux, Conor Joseph Meehan, Philip Supply

## Abstract

The human- and animal-adapted lineages of the *Mycobacterium tuberculosis complex* (MTBC) are thought to have clonally expanded from a common progenitor in Africa. However, the molecular events that accompanied this emergence remain largely unknown. Here, we describe two MTBC strains isolated from patients with multidrug-resistant tuberculosis, representing an as-yet-unknown lineage, named Lineage 8 (L8), seemingly restricted to the African Great Lakes region. Using genome-based phylogenetic reconstruction, we show that L8 is a sister clade to the known MTBC lineages. Comparison with other complete mycobacterial genomes indicate that the divergence of L8 preceded the loss of the *cobF* genome region - involved in the cobalamin/vitamin B12 synthesis - and gene interruptions in a subsequent common ancestor shared by all other known MTBC lineages. This discovery further supports an East African origin for the MTBC and provides additional molecular clues on the ancestral genome reduction associated with adaptation to a pathogenic lifestyle.

## Introduction

Tuberculosis (TB), caused by members of the *Mycobacterium tuberculosis complex* (MTBC), is among the ancient scourges of humankind^1^, and remains the leading cause of mortality globally due to an infectious disease^2^. Intense research has been dedicated to decipher the evolutionary history of the MTBC and to understand the causes underlying the worldwide spread of TB^3–5^. Current genome data show that the MTBC is comprised of the five human-adapted lineages representing *M. tuberculosis sensu stricto* (L1–4, and L7), two other human-adapted lineages traditionally referred to as *M. africanum* (L5-6) and at least nine animal-adapted lineages^6^. Africa is the only continent where all MTBC lineages are present, suggesting that the MTBC emerged from a common ancestor therein and then clonally expanded to the rest of the world following human migrations^3,7–10^. However, the genomic traits of this common ancestor and the region from which this expansion took place in Africa remain unknown. Whole genome sequencing (WGS) analyses showed that rare human TB bacilli with a smooth colony morphotype, highly restricted to the Horn of Africa and named *Mycobacterium canettii* (alias smooth tubercle bacilli or STB) represent early evolutionary branching lineages that predate the emergence of the most recent common ancestor (MRCA) of the MTBC (or of the rest of the MTBC, if *M. canettii* is considered to be part of the complex)^4,11,12^ Indeed, whereas known MTBC strains differ by no more than ~2,000 Single Nucleotide Polymorphisms (SNPs)^13^, *M. canettii* strains are 10 to 25-fold more genetically diverse and separated by at least 14,000 SNPs from the hitherto known MTBC MRCA^4,12^. Moreover, *M. canettii* strains are less virulent and possess highly mosaic genomes, possibly reflecting adaptation to an environmental reservoir favouring active lateral gene flow^4,14–16^. These biological differences support the existence of lineages that reflect intermediate stages in the evolution from *M. canettii* towards the obligate MTBC pathogens.

Here, we describe two exceptional strains representing a new clade, diverging before the MRCA of the other MTBC lineages. These two strains were discovered in two independent analyses, and were both multidrug-resistant (MDR; i.e. resistant to at least rifampicin and isoniazid). One was isolated from a TB patient in Rwanda through an ongoing MDR-TB diagnostic trial in Africa. The second isolate was from a patient in Uganda, and was discovered upon screening publicly available draft genome datasets, where it was misclassified as an *M. bovis* strain. We used PacBio and Illumina WGS to reconstruct the full circular genome of the Rwandan strain. We used these data and the available Illumina sequencing data of the Ugandan strain to reconstitute the phylogeny of this novel lineage, which we named Lineage 8 (L8), and further investigate molecular and evolutionary events associated with the emergence of the MTBC.

## Results

### Patient with the L8 TB strain in Rwanda

The strain in Rwanda was isolated from a male patient, aged 77 years, HIV-negative, resident of Rulindo district bordering with the Southwest of Uganda, and who had lived in Uganda previously. The patient was diagnosed with rifampicin-resistant TB based on standard Xpert MTB/RIF testing (Xpert; Cepheid, Sunnyvale, CA, USA), which probes for mutations in the rifampicin resistance-determining region of the *rpoB* gene of MTBC strains^17^. The results of the assay showed a rare delayed probe B reaction (~3% prevalence in Rwanda)^18^, presumed (and later confirmed; see below) to be due to the rifampicin resistance-associated Asp435Tyr mutation^19^.

Per routine practice, the patient was initiated on standard short-course MDR-TB treatment (i.e. 9-month WHO-endorsed MDR-TB regimen, including moxifloxacin, kanamycin, protionamide, ethambutol, clofazimine, high dose isoniazid and pyrazinamide)^20^. However, the patient developed hypotension, and eventually died due to probable cardiac failure, after 20 days of treatment. Phenotypic drug-susceptibility testing (DST) confirmed resistance to both rifampicin and isoniazid, and susceptibility to other anti-TB drugs including ethambutol, fluoroquinolones, and second-line injectables.

### Growth characteristics and biochemical properties of the Rwandan strain

The strain from Rwanda was grown in 12.5 days on Mycobacterial Growth Indicator Tubes. Colonies were observed on the fifth week after initial inoculation on Löwenstein-Jensen medium, indicating a slow grower phenotype with rough colonies (Figure 1). The strain also displayed archetypal biochemical characteristics of *M. tuberculosis sensu stricto*, including niacin production combined with urease hydrolysis (Table 1).

**Table 1.**
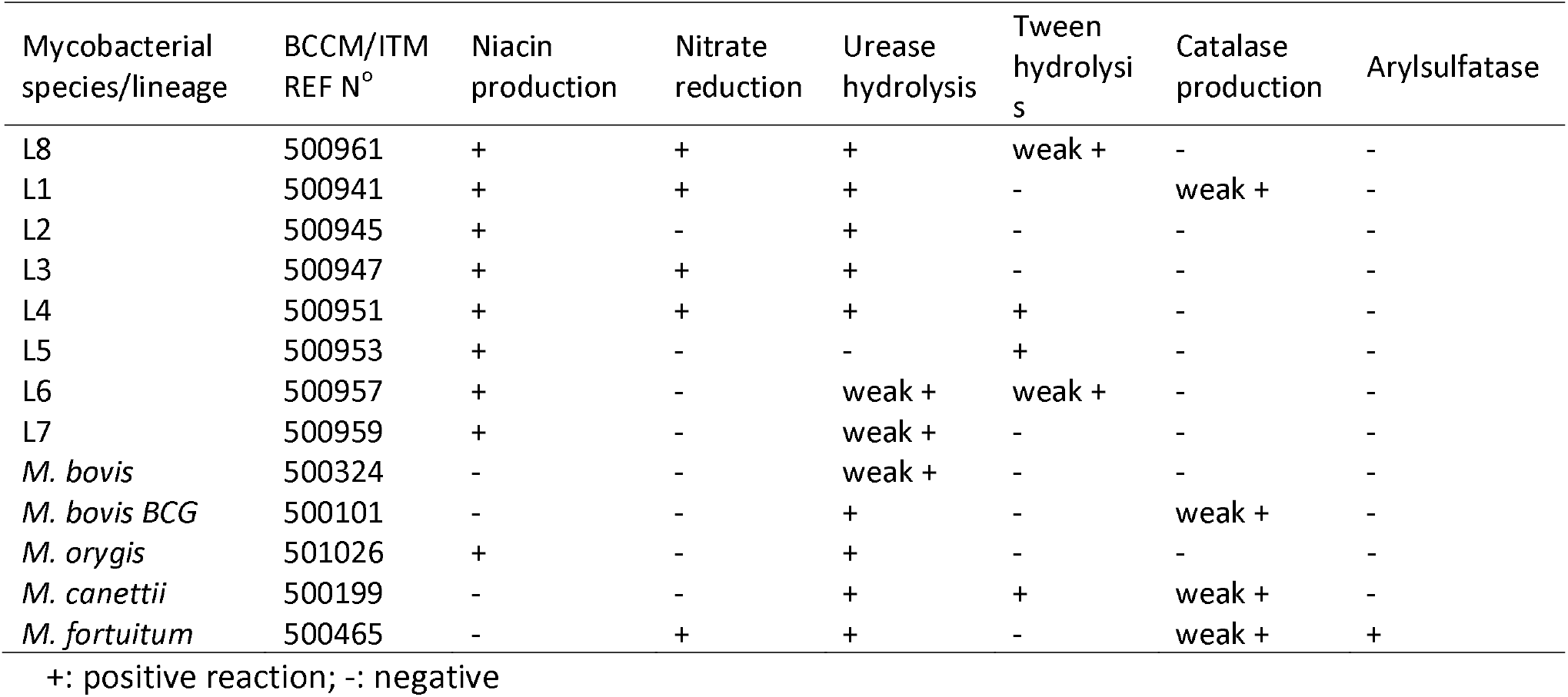
Standard biochemical characteristics of selected mycobacterial species or *M. tuberculosis* complex lineages/subspecies versus L8

| Mycobacterial species/lineage | BCCM/ITM REF N° | Niacin production | Nitrate reduction | Urease hydrolysis | Tween hydrolysis | Catalase production | Arylsulfatase |
| --- | --- | --- | --- | --- | --- | --- | --- |
| L8 | 500961 | + | + | + | weak + | - | - |
| L1 | 500941 | + | + | + | - | weak + | - |
| L2 | 500945 | + | - | + | - | - | - |
| L3 | 500947 | + | + | + | - | - | - |
| L4 | 500951 | + | + | + | + | - | - |
| L5 | 500953 | + | - | - | + | - | - |
| L6 | 500957 | + | - | weak + | weak + | - | - |
| L7 | 500959 | + | - | weak + | - | - | - |
| *M. bovis* | 500324 | - | - | weak + | - | - | - |
| *M. bovis BCG* | 500101 | - | - | + | - | weak + | - |
| *M. orygis* | 501026 | + | - | + | - | - | - |
| *M. canettii* | 500199 | - | - | + | + | weak + | - |
| *M. fortuitum* | 500465 | - | + | + | - | weak + | + |

+: positive reaction; -: negative

**Figure 1.**
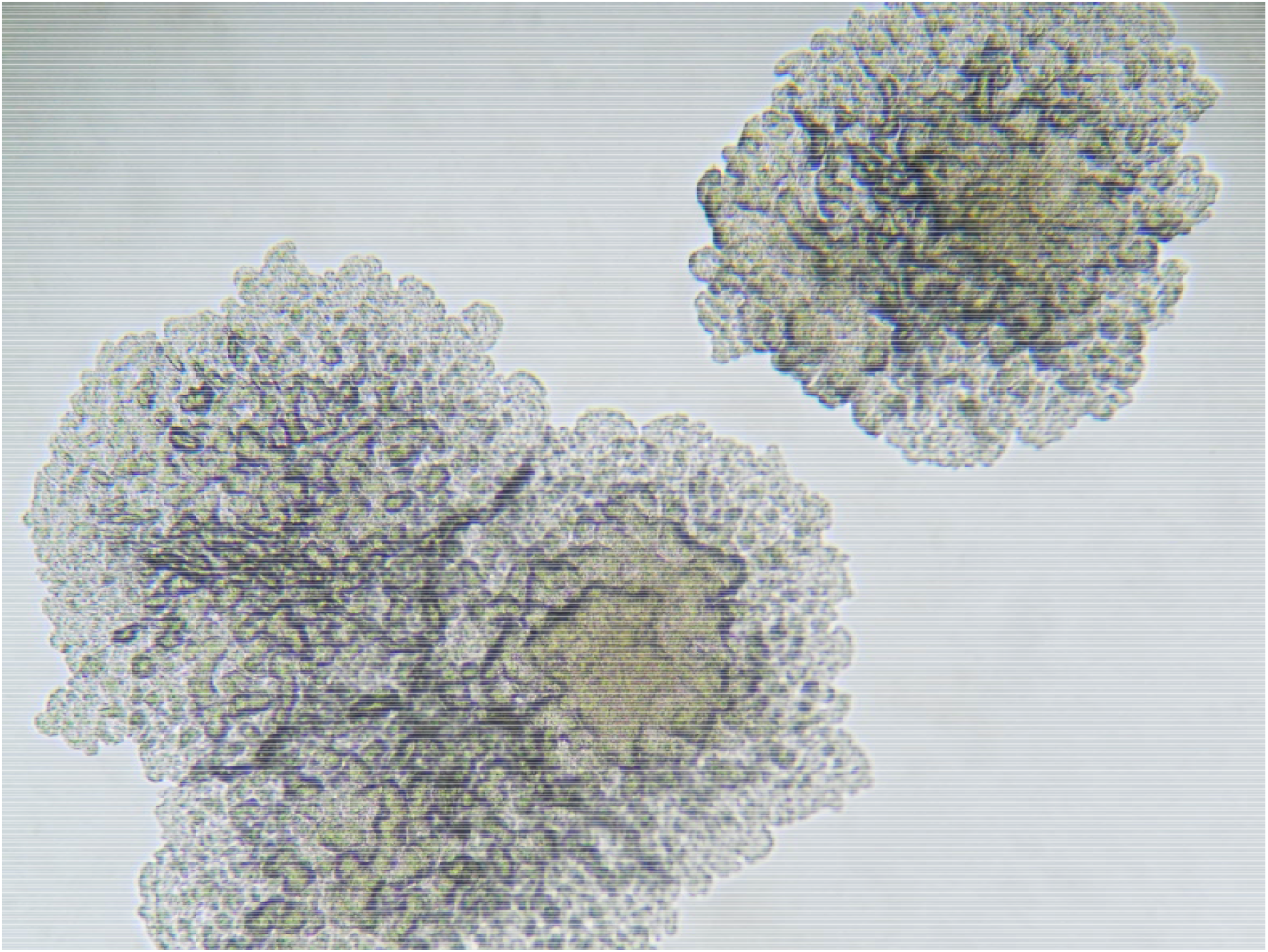
Microscopic image of L8 on Dubos agar medium showing typical rough colonies (read at 100x).

### Genotypic resistance and SNP profile of the Rwandan strain by Deeplex Myc-TB

Following the MDR-TB diagnosis, the strain was included in the first set of tests for an ongoing MDR-TB diagnostic trial “DIAgnostics for MDR-TB in Africa (DIAMA) Clinicaltrials.gov, NCT03303963”, evaluating a new targeted deep sequencing assay, called Deeplex-MycTB. Deeplex-MycTB testing confirmed the presence of the *rpoB* Asp435Tyr mutation conferring rifampicin resistance, along with the *inhA* Ser94Ala mutation conferring isoniazid resistance, consistently with the MDR phenotype identified by phenotypic DST (**Figure 2**). This strain also harboured two alleles in phylogenetic positions in *embB* (Ala378) and *gidB* (Ala205) not associated with resistance to ethambutol or streptomycin, which were both shared by several MTBC lineages (L1, 5, 6, 7, and animal lineages) and *M. canettii*^21^. In addition, nine other - so far uncharacterized - SNPs were identified in six of the 18 gene targets interrogated by the assay (**Figure 2**). Moreover, this test detected an atypical spoligotype pattern, 1111100000000000000000000000000001110000000 (**Figure 2**), which was further confirmed by conventional membrane-based spoligotyping testing. This spoligotype pattern was unique in the global spoligotype database that comprises 111,637 MTBC isolates from 131 countries^22^.

**Figure 2.**
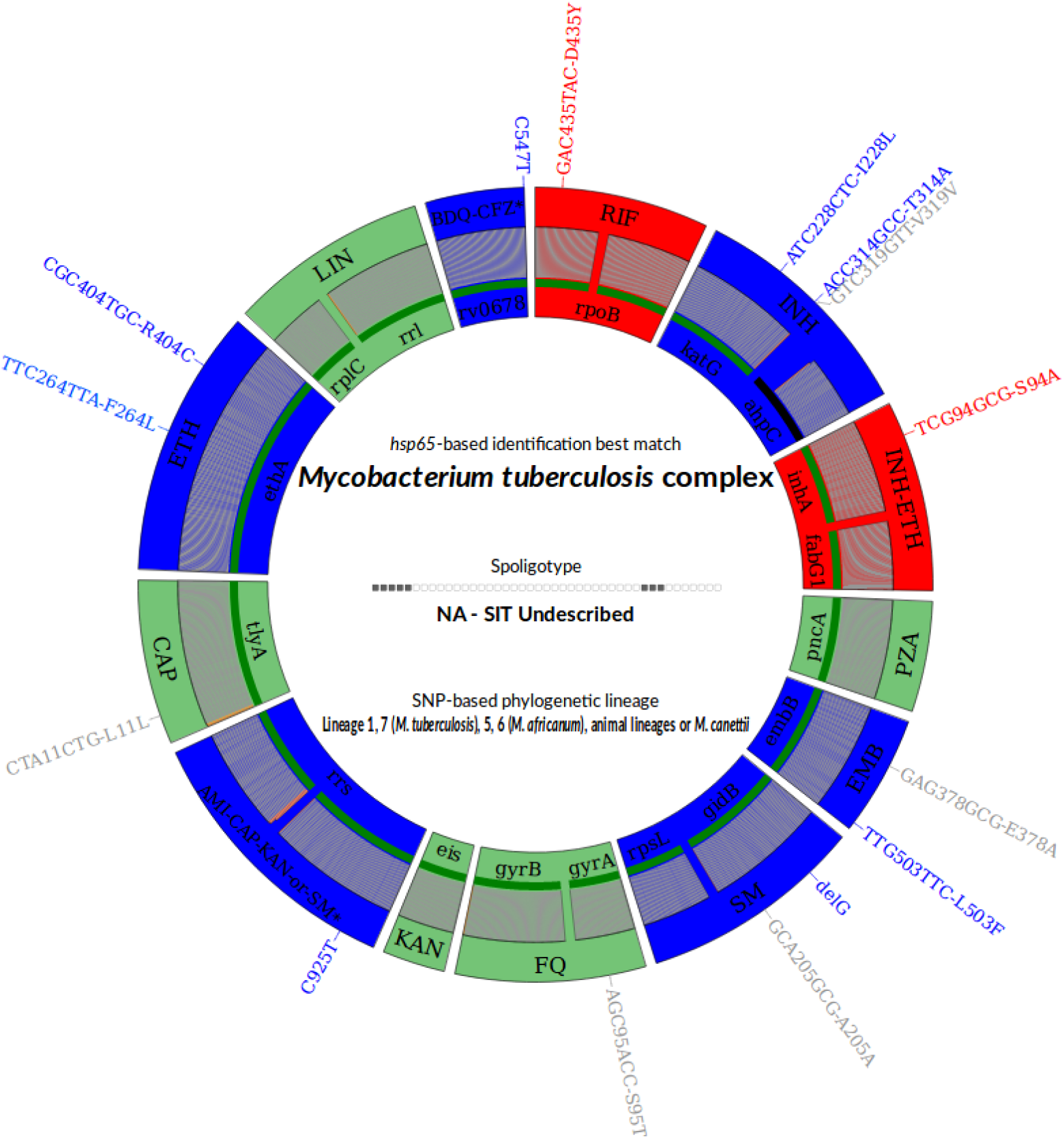
Deeplex-MycTB results identifying a multidrug resistant strain from Rwanda with an atypical genotypic profile in the *M. tuberculosis* complex. Target gene regions are grouped within sectors in a circular map according to the tuberculous drug resistance with which they are associated. The two sectors in red indicate regions where rifampicin and isoniazid resistance associated mutations are detected. The multiple sectors in blue refer to regions where as yet uncharacterized mutations are detected, while sectors in green indicate regions where no mutation or only mutations not associated with resistance (shown in gray around the map) were detected. Green lines above gene names represent the reference sequences with coverage breadth above 95%. Limits of detection (LOD) of potential heteroresistance (reflected by subpopulations of reads bearing a mutation), depending on the coverage depths over target sequence positions, are represented by grey (LOD 3%) and orange zones (variable LOD >3%–80%; only seen in extremities of a few targets, such as the two rrs regions) above the reference sequences within the sectors. Information on an unrecognized spoligotype, an equivocal SNP-based and on mycobacterial species identification, based on *hsp65* sequence best match, are shown in the centre of the circle. *AMI, amikacin; BDQ, bedaquiline; CAP, capreomycin; CFZ, clofazimine; EMB, ethambutol; ETH, ethionamide; FQ, fluoroquinolones; KAN, kanamycin; LIN, linezolid; INH, isoniazid; PZA, pyrazinamide; RIF, rifampin; SM, streptomycin; SIT, spoligotype international type.

### WGS analysis and phylogenetic position of the Rwandan and Ugandan strains

Results from WGS analysis of the Rwandan strain using Illumina sequencing confirmed all Deeplex-MycTB findings.

The strain isolated in Uganda was discovered independently upon screening global MTBC genome datasets publicly available on NCBI/EBI, comprising approximately 20,000 genomes in total. From subsequent processing with our WGS analysis pipeline, we found 1 genome that did not classify in any of the 7 human-adapted lineages or 9 animal-adapted ecotypes known at the time, but had been misclassified as *M. bovis* isolated from a human patient^23^. These WGS data revealed a similar spoligotype 1111100000000000000000000000000001111000111, characterized by the presence of spacers 1 to 5 and 34 to 37 (vs 34-36 in the Rwandan strain) with all intervening spacers missing. Moreover, the Ugandan strain also shared the same *rpoB* Asp435Tyr and *inhA* Ser94Ala mutations and the same sequence alleles in *embB* and *gidB*. The Ugandan strain contained an additional *katG* Ser315Thr mutation conferring isoniazid resistance, as well as the *embA* C-11A and *embB* Asp328Tyr mutations, and two *pncA* missense mutations, predictive of pyrazinamide resistance. Moreover, only three of the nine aforementioned uncharacterized SNPs detected by Deeplex-MycTB were shared between both strains. To further assess the relationships between both strains and in comparison to other MTBC strains, a maximum likelihood phylogeny was inferred from 241 MTBC genomes, including representatives of all known human- and animal-adapted lineages^6^ and using a *M. canettii* strain as an outgroup. *M. canettii* represents the closest outgroup to the MTBC including L8, as shown by subsequent comparative analysis of a complete L8 genome (see below), and previous observations of ~2.0 Mb larger genomes and substantially lower average nucleotide identities of phylogenetically closest non-tuberculous mycobacterial species such as *M. marinum* and *M. kansasii*^4,24,25^. This reconstruction revealed a unique phylogenetic position of the two new genomes from Rwanda and Uganda (**Figure 3**), representing a newly characterised monophyletic clade in which none of the known MTBC genomes are contained. A core genome-based phylogeny was also constructed from representatives of the MTBC lineages as well as *M. canettii*, *M. marinum* and *M. kansasii* (Supplementary figure 1). This phylogeny confirmed the placement of the L8 Rwandan strain as being a new clade between *M. canettii* and the other lineages of the MTBC. Based on these phylogenies, this clade shares a MRCA with the rest of the MTBC, thus representing a new sister clade to the known MTBC, which we named Lineage 8 (L8). Comprehensive SNP analysis identified a total of 189 SNPs separating both L8 genomes, which is within the range of zero to 700 SNPs found between any two strains within any of the lineages 1 to 7 of the MTBC^13^. On average, two strains of the MTBC including L8 differed by 1443 SNPs (corresponding to 0.04% of the genome, excluding repetitive/‘problematic’ regions), thus at least an order of magnitude lower than the SNP distance separating any MTBC strain from *M. canettii*^4,13^.

**Figure 3.**
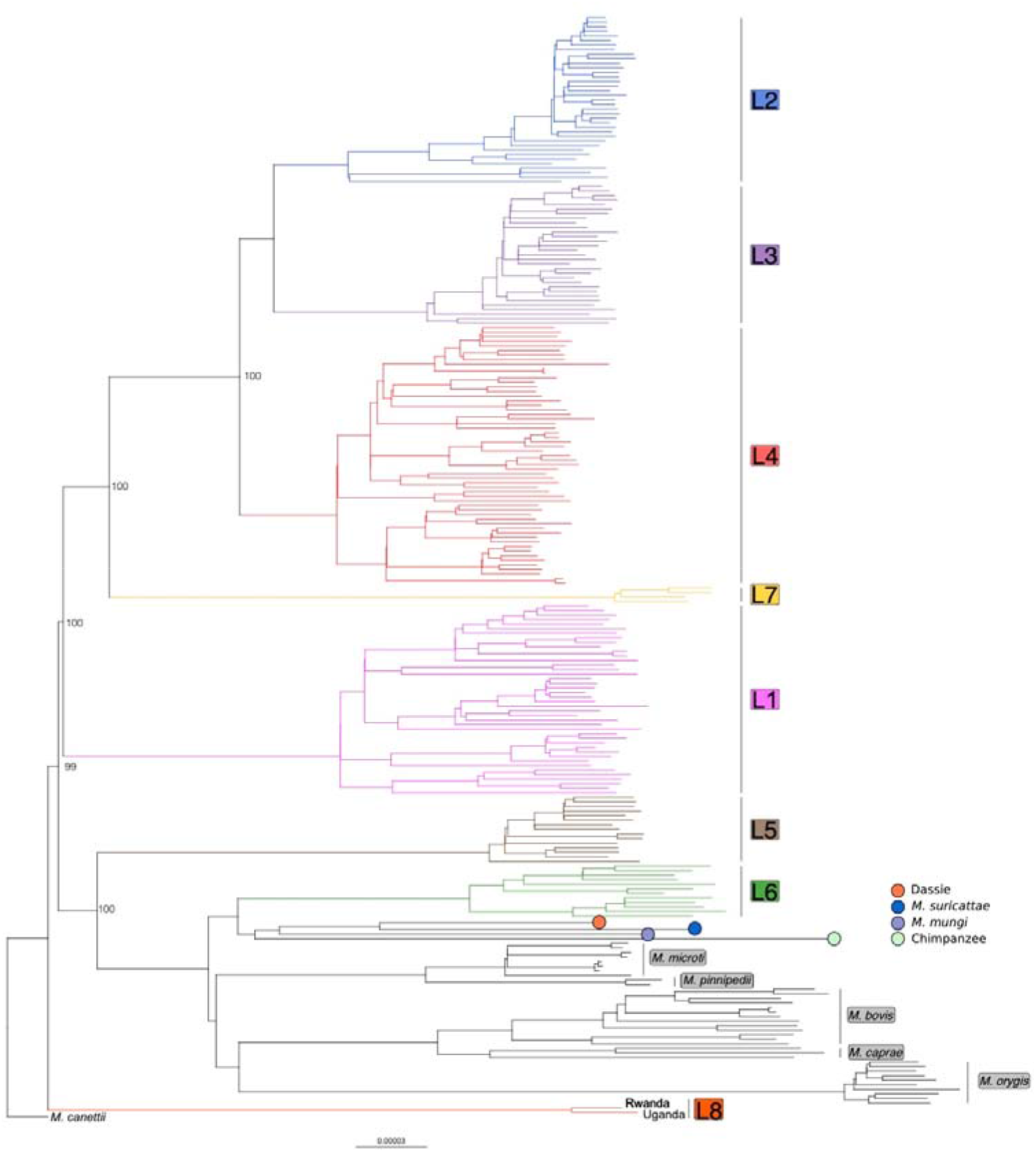
Maximum likelihood phylogeny of 241 MTBC genomes, inferred from 43,442 variable positions. The scale bar indicates the number of substitutions per polymorphic site. Branches corresponding to human-adapted strains are coloured and branches corresponding to animal-adapted strains are depicted in black. The phylogeny is rooted on *M. canettii* and bootstrap values are shown for the most important splits.

The absence of any matching pattern in the global spoligotype database, as well as the lack of detection of this clade in previous large WGS datasets of MTBC strains from global sources, suggests that L8 is rare and seemingly geographically restricted to the African Great Lakes region. Specifically, the L8 spoligotype signature and the 3 SNPs shared by both L8 strains were not detected in any of 115 MTBC genomes from a previous drug resistance survey in Uganda^26^, nor in 380 rifampicin-resistant strains from Rwanda collected between 1991 and 2018, from routine drug resistance surveillance as well as various drug resistance surveys^27–29^. Furthermore, among 14 other available isolates out of 27 from Uganda and Rwanda tested by Gene Xpert MTB/RIF that showed the same delayed probe B as L8, none displayed the L8 signatures when tested by Deeplex Myc-TB or by classical spoligotyping. Likewise, none of > 1,500 clinical samples from TB patients tested by Deeplex-MycTB from a recent nationwide drug resistance survey performed in the Democratic Republic of Congo displayed the L8 spoligotype signature or the specific SNPs.

### Defining features of a complete L8 genome

To further assess the sister position of L8 and split from the remaining MTBC inferred from SNP analysis, the Rwandan strain was subjected to long read-based PacBio sequencing. Comparison of the obtained assembly with 36 available complete genomes of MTBC members, comprising L1-L4 (including H37Rv), *M. africanum* (L6) and *M. bovis* strains, showed a highly syntenic organization, with no major structural rearrangement between both groups. Although the assembled L8 genome of 4,379,493 bp was within the 4.34-4.43 Mb size range of the other MTBC genomes, it was 30 kb smaller than the 4.41-Mb mean size of genomes of *M. tuberculosis sensu stricto*^30^. However, the largest part of this gap was accounted by the absence of three genomic regions in L8, corresponding to regions of difference (RDs) known to be variably present or absent in other MTBC (sub)lineages^31,32^(Supplementary Table 2). These include a 9.3-kb PhiRv1 prophage region (RD3), as well as 10.0-kb and 8.5-kb segments corresponding to RD14 and RD5, comprising the *plcABC* gene cluster and the *plcD* gene regions, respectively^31^. In L8, each of the two latter regions only contained one copy of the IS*6110* insertion sequence, devoid of direct repeats (DRs) that normally flank IS*6110* copies after transposition, indicating that these deletions in L8 resulted from recombination between two adjacent IS*6110* copies with loss of the intervening sequences^33^. These mobile DNA-related deletions, which also arose independently in several other MTBC branches^31,34^, probably occurred after the divergence of L8 from the other MTBC lineages. Apart from these three deletions and two dozen repetitive/multicopy genes (IS6110-related, PE/PPE-, or Mce-encoding), we only found 5 non-repetitive genes, included in two small segments (3.4 and 4.4 kb), which were undetected in the complete L8 genome while being present in reference MTBC genomes (Supplementary Table 2).

Conversely, a particular 4.4 kb genome region was present in the genomes of both L8 strains and in *M. canettii*, but absent in the 36 available complete genomes of MTBC members (Supplementary Table 3). This region comprises the cobF gene (Figure 4), encoding the precorrin 6A synthase involved in the cobalamin/vitamin B12 synthesis, along with two other genes, respectively encoding a PE-PGRS protein family member and a protein of unknown function. This region shared by the L8 and the *M. canettii* genomes is also present in the phylogenetically proximal non-tuberculous mycobacterial species *M. marinum* and *M. kansasii* (Supplementary Table 3). Via BLAST analysis, we further confirmed the systematic absence of *cobF* in any of 6,456 quality draft genome assemblies available as of January 2020 from the NCBI, from strains belonging to lineages 1 to 7 or the animal lineages of the MTBC. Moreover, we thereby determined that the junction between the sequences flanking the cobF deletion was at the same nucleotide position in all but 6 of these genomes, resulting in the truncation of *rv0943c* and *rv0944* genes as seen in the complete MTBC genomes (Fig. 4). Consistent with the clonal evolution of the MTBC with negligible, if any, horizontal gene transfer between strains^1,14,32,35^ this perfect conservation of this sequence junction suggests that cobF was lost in the MRCA of the other MTBC lineages, after its divergence from L8. The 6 exceptions were 3 strains from lineage 4.3 and 3 from lineage 3 that showed slightly larger deletions, including the 5’ region of *rv0943c* or the 5’ region of *rv0943c*, *rv0944* and the 5’ region of *rv0945*, respectively, suggesting probable subsequent deletion events in particular sub-branches of these lineages.

**Figure 4.**
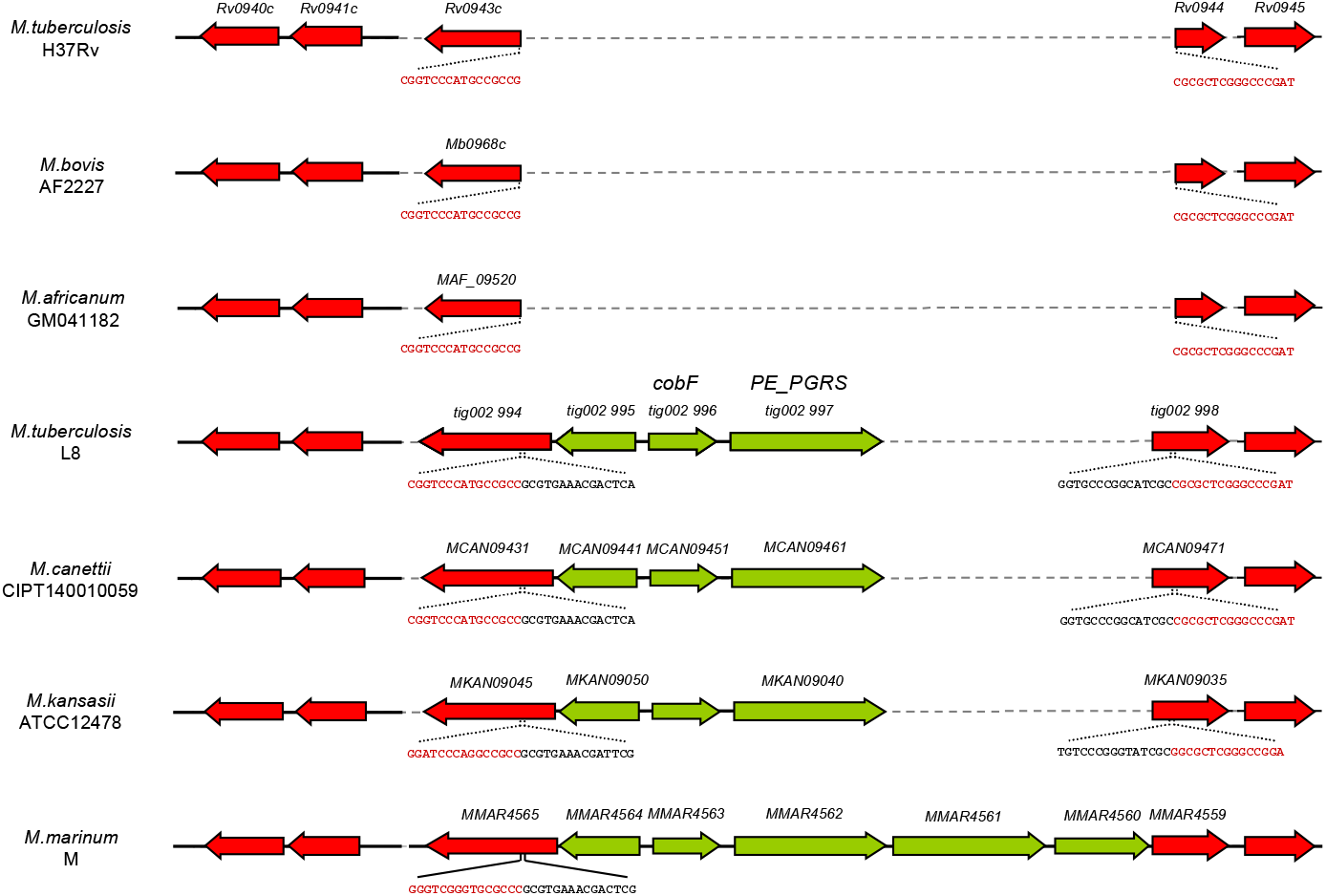
Aligned genome segments showing the *cobF* gene region in *M. tuberculosis* L8, *M. canettii* CIPT140010059 (alias STB-A), M. kansasii ATCC12478 and M. marinum M strains, and the corresponding deletion in *M. tuberculosis* H37Rv, M. bovis AF2122/97, and *M. africanum* GM041182. Coding sequences of this region are shown in green, and flanking coding sequences in red. Sequences flanking the deletion point in truncated genes in *M. tuberculosis*, *M. africanum* and *M. bovis*, and in the cobF region present in *M. canettii*, *M. kansasii* and *M. marinum* are indicated in red and black, respectively. Dashed lines correspond to missing segment parts relatively to the longest segment found in *M. marinum*.

In contrast, none of the almost 900 other genes specifically identified in the *M. canettii* genomes, and absent in the other MTBC genomes, were found in the L8 genome. The latter finding thus supports the close relationship with the previously known MTBC lineages indicated by the SNP-based phylogeny, as well as the outgroup position of *M. canettii* relatively to the MTBC including L8.

Further evidence for the early branching of L8 relative to the rest of the MTBC comes from examination of interrupted coding sequences (ICDSs). These ICDSs correspond to frameshifts or in-frame stop codons detected in genes originally intact in a common progenitor, thus putatively representing so-called molecular scars inherited during progressive pseudogenization of the MTBC genomes^36,37^. Four orthologues of MTBC ICDSs were previously found to be intact in the genomes of *M. canettii* strains, as well as in *M. marinum* and *M. kansasii* ^4^. One of these four orthologues (*pks8*), which belongs to a multigene family encoding polyketide synthases involved in the biosynthesis of important cell envelope lipids^38^, was also intact in the genomes of both L8 strains (Figure 5 and Supplementary Table 4). Moreover, we found an additional orthologue of MTBC ICDSs (i.e. *rv3899c-rv3900c*), coding for a conserved hypothetical protein, which was intact in the genomes of *M. canettii*, *M. kansasii*, *M. marinum* and both L8 strains (Supplementary Table 4). These two molecular scars were also likely acquired by the other MTBC lineages after their divergence from the common progenitor shared with L8.

**Figure 5.**
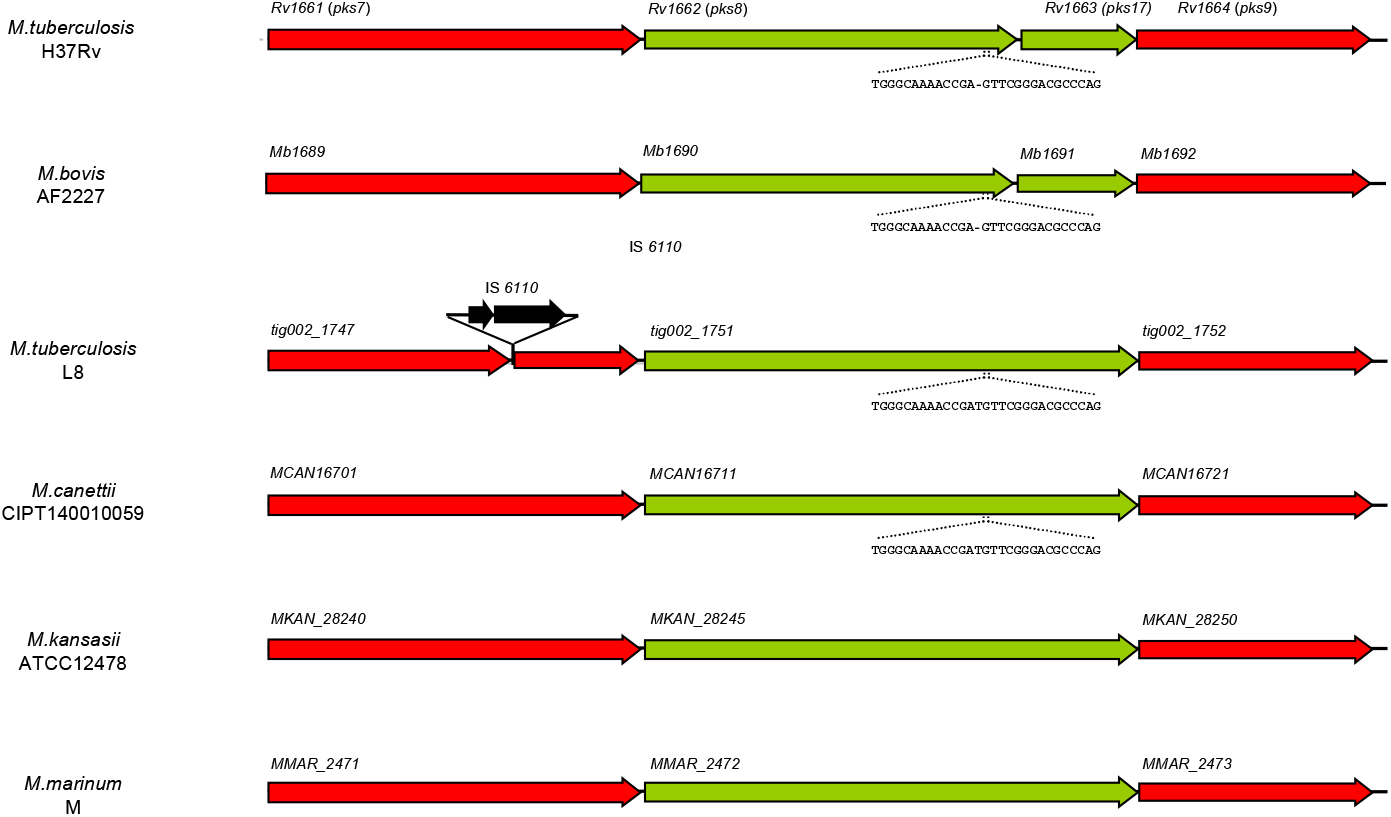
Aligned genome segments showing the interrupted coding sequences *pks8/17* in *M. tuberculosis* H37Rv and *M. bovis* AF2122/97 gene region, and complete *pks8* genes in L8, *M. canettii* CIPT140010059 (alias STB-A) and *M. kansasii* ATCC12478. *pks8/17* and *pks8* coding sequences are shown in green, and flanking genes in red. Sequences flanking the 1-nucleotide deletion and resulting in a frameshift in *M. tuberculosis* complex strains are indicated. Dashed lines correspond to missing segment parts relatively to the longest segment found in *M. canettii*.

The assembled L8 genome also included 48 out of 50 genes (the exceptions are rv3513c encoding the probable fatty-Acid-Coa ligase FadD18 and the PhiRv1 region; see above) present in MTBC members but not found in any of the *M. canettii* genomes, including a number of genes putatively acquired through horizontal gene transfer by the common ancestor of the MTBC after its separation from *M. canettii*^4^ (Supplementary Table 5). This observation additionally supports both the close relationship with the previously known MTBC lineages and the outgroup position of *M. canettii* relatively to the MTBC including L8. Likewise, consistent with the rough colony morphotype of the Rwandan strain, both L8 strains displayed the single polyketide-synthase-encoding *pks5* gene configuration shared by all MTBC members, instead of the dual *pks5* conformation found in *M. canettii* strains involved in the smooth colony phenotype of the latter strains^15^. Thus, the recombination between the two *pks5* genes and the loss of the intervening *pap* gene, thought to have resulted in surface remodelling and incremental gain of virulence after the phylogenetic separation from *M. canettii*^15^, already existed in the common progenitor of L8 and the rest of the MTBC. Moreover, both L8 strains also contained the intact TbD1 and RD9 regions, shared by the other “ancestral” *M. tuberculosis* lineages (L1, L7) but subsequently lost by the so-called “modern” lineages of *M. tuberculosis* (TbD1 lost in L2-4), *M. africanum* (L5 and L6) and the animal lineages (RD9)^31^.

In contrast to the highly clonal population structure of the MTBC, *M. canettii* strains are highly recombinogenic, as apparent from mosaic sequence arrangements in their genomes and functional DNA transfer between *M. canettii* strains mediated by a distributive conjugal transfer (DCT)-like mechanism^4,39^. However, no significant genome-wide recombination signal was detected by ClonalFrameML analysis^40^ between L8 and other MTBC strains (Supplementary Figure 2). In particular, and in contrast to the numerous recombination segments in *M. canettii*^4^, the complete L8 genome only contained 26 possible recombination segments, yet the longest of these was 607bp and the average length was 142bp.

## Discussion

The discovery of L8 provides unique insights into an ancestor of the MTBC that existed after the *pks5*-recombination-mediated surface remodelling, which occurred after separation of the MTBC MRCA from the *M. canettii* clade, but preceded the loss of the *cobF* region and gene interruptions in a later common ancestor of the other MTBC lineages. The seeming restriction of this lineage to the African Great Lakes region represents new evidence supporting an origin for the MTBC in the eastern part of the African continent. These findings reinforce results from previous work suggesting an East-rather than a West African origin of the MTBC^3,4,9,11,41^.

A distinct ecological niche, linked to a potential environmental reservoir, has been hypothesized to explain the marked geographic restriction of *M. canettii* strains to the Horn of Africa, the lower persistence of these strains in infection models as well as their genome mosaicism implying multiple DNA recombination events within the *M. canettii* strain pool^4,12^. However, although our analysis is limited to two genomes identified to date, our results suggest that L8 is as clonal as the rest of the MTBC^3,14,34,42^. Moreover, the observation that both L8 strains share two uncommon rifampicin- and isoniazid-resistance conferring mutations in *rpoB* and *inhA* suggests that multidrug resistance was already acquired in the common ancestor of these two strains. Isoniazid and rifampicin were introduced in TB treatments in the African Great Lakes region in the late fifties and 1983, respectively (Dr. Armand Van Deun, personal communication). These shared MDR-defining mutations, and the detection of these isolates in human patients in both cases (with reported absence of previous TB history for the Rwandan patient), suggest that these patients were infected with an already-resistant strain, which was exposed to drug selective pressure already decades ago and had been likely circulating in the community for some time. Overall, this pattern thus suggests human-to-human transmission rather than infection from a non-human source. While based on only two initial strains, these results are consistent with the presumed scenario of a human rather than a zoonotic origin for the MTBC^31,43^.

Of note, given the above timeline of introduction of rifampicin and isoniazid in both countries, the ~100 SNPs distance separating these two strains from their MRCA would imply a rapid molecular clock for L8, in the range of the upper bound of 2.2 SNPs/genome/year most recently estimated for other MTBC clades^44^. However, this mutation rate cannot be confirmed until additional L8 samples are uncovered.

Remarkably, the absence of other L8 strains in datasets from Uganda, Rwanda and DRC, together comprising more than 2,000 strains, suggests that L8 is rare even within the African Great Lakes region. Such scarcity is compatible with selective sweeps of later branching MTBC strains, introduced more recently into the region. Similar scenarios have also been proposed to explain the slow apparent replacement of MTBC L5 and L6 by L4 in West Africa^45–47^ and the restriction of L7 to Northern Ethiopia^48^.

Loss-of-function linked to the deletion of *cobF* is a plausible candidate molecular event involved in such a replacement scenario for L8. Indeed, loss-of-function appears to be an important mechanism driving the pathoadaptive evolution of the TB pathogen, as shown for the role of the loss of lipo-oligosaccharide production (via recombination in the *pks5* locus)^15^ in the evolution towards increased virulence from *M. canettii* to MTBC strains. Likewise, loss of secretion of PPE-MPTR and PE_PGRS proteins by the type VII secretion system ESX-5 (via mutations of the ppe38 locus) has been involved in the hypervirulence of recent branches of L2 (alias “modern” Beijing) strains^49^. The loss of the *cobF* region in the other MTBC lineages, inferred from comparative genomics with *M. canettii* and non-tuberculous mycobacteria^4^, was previously hypothesised to reflect enhanced adaptation to an intracellular parasitic lifestyle^50^. Indeed, the cobalamin/vitamin B12 synthesis pathway, of which the *cobF*-encoded precorring-6a synthase is a component, represents a highly complex and energy consuming process with about 30 enzymatic steps^51^. While the absence of this component may not entirely ablate cobalamin biosynthesis^52,53^, its loss might have resulted in gain of fitness and reflect enhanced pathogenic professionalisation, by economical reliance upon the mammalian host environment as source of vitamin B12. As an additional plausible but not necessarily mutually exclusive hypothesis, such selective sweep of L8 might have been (further) enhanced by the loss of theTbD1 region in later branching MTBC strains. This region, which we also found intact in L8, as is the case in the “ancestral” *M. tuberculosis* lineages L1 and L7, encodes members of the mycobacterial membrane protein families MmpL. Very recent findings indicate that the loss of this region in later branching MTBC strains was also associated with a gain of virulence, and the deletion of TbD1 at the origin of the “modern” *M. tuberculosis* lineages L2/L3/L4 has therefore been suggested as a key driver for their global epidemic spread^54^. If true, more recently emerged or introduced *cobF*- and TbD1-deleted strains might conceivably have largely outcompeted L8 strains. This hypothesis could be explored by assessing the growth and the virulence/fitness in cellular and animal models, of recombinant *cobF*- and/or TbD1-knock-out, as well as *cobF*- and/or TbD1-knock-in strains, derived from the available Rwandan L8 strain and other MTBC strains, respectively.

In conclusion, our genomic data, on an as-yet-unknown ancestral stage between the MTBC and the putative progenitor pool of *M. canettii*-like mycobacteria, thus suggest further experiments to examine candidate molecular events potentially involved in the pathoadaptive evolution of *M. tuberculosis*. The discovery of such rare strains raises the possibility for the existence of further extant strains, especially in Eastern Africa, representing other clades further closing the biological gap between the MTBC and *M. canettii*.

## Methods

### Rwandan patient recruitment and ethics statement

The patient in Rwanda was recruited into, and gave informed consent for, the DIAMA study which was approved by the Rwanda National Ethical committee (IRB 00001497 of IORG0001100; Ref No.0069/RNEC/2017).

### Phenotypic characterization

We studied conventional mycobacterial growth and biochemical characteristics including colony morphology, niacin production, nitrate reduction, p-nitro benzoic acid growth inhibition, catalase production, urea hydrolysis, tween 80 hydrolysis, and thiophene carboxylic acid hydrazide growth inhibition^55^. For comparative purpose, a reference set of the seven known human-adapted MTBC lineages^56^, together with *M. canettii* (BCCM/ITM500199), *M. bovis* (BCCM/ITM500324), *M. bovis* BCG (BCCM/ITM500101), and *M. orygis* (BCCM/ITM501026) strains were processed with the novel strain isolated in Rwanda. Moreover, phenotypic drug-susceptibility testing to first- and second-line anti-TB drugs was done using the proportion method^57^. The strain isolated in Uganda was not available for phenotypic characterization.

### Targeted and whole genome sequencing of the Rwandan strain

Targeted sequencing was performed by using the Deeplex-MycTB assay^58^ (Genoscreen, France). Briefly, this assay relies on a 24-plexed PCR amplification of mycobacterial species identification (*hsp65*), genotyping (spoligotyping and phylogenetic single nucleotide polymorphisms (SNPs)) and 18 *M. tuberculosis* complex drug resistance-associated gene targets. This test and short-read Illumina-based WGS were performed on the Rwandan strain as follows. A bead beating method was used to extract DNA from colonies (Supplementary method 1). Libraries of Deeplex-MycTB amplicons or genome fragments were constructed using the Nextera XT kit and sequenced on an Illumina MiSeq platform with paired end, 150-bp read lengths (Illumina, CA, USA). DNA extraction suitable for PacBio SMRT sequencing was performed using the Genomic DNA Buffer Set (Qiagen Inc, Germantown, Maryland, USA) (Supplementary method 2). Sequencing was performed on a PacBio RS II using the SMRT technology.

### Deeplex-MycTB analysis and spoligotyping

Analysis of the Deeplex-MycTB sequencing data, including SNP calling and spoligotype identification, was performed by read mapping on *M. tuberculosis* H37Rv sequence references, using a parameterized web application (GenoScreen)^58^. Membrane-based spoligotyping was performed as described previously^59^.

### Illumina whole genome sequencing analysis

Raw genomic reads from the newly sequenced L8 genome from Rwanda and the L8 genome from Uganda (SAMN02567762) were processed as previously described^60^. Briefly, the reads were trimmed with Trimmomatic v0.33.22^61^ and reads larger than 20 bp were kept. The software SeqPrep (https://github.com/jstjohn/SeqPrep) was used to identify and merge any overlapping paired-end reads. The resulting reads were aligned to the reconstructed ancestral sequence of the MTBC^62^ using the mem algorithm of BWA v0.7.13^63^. Duplicated reads were marked using the MarkDuplicates module of Picard v2.9.1 (https://github.com/broadinstitute/picard) and local realignment of reads around InDels was performed using the RealignerTargetCreator and IndelRealigner modules of GATK v3.4.0^64^. SNPs were called with Samtools v1.2 mpileup^65^ and VarScan v2.4.1^66^ using the following thresholds: minimum mapping quality of 20, minimum base quality at a position of 20, minimum read depth at a position of 7X, maximum strand bias for a position 90%. The spoligotype pattern of the strain from Uganda was extracted *in silico* from the raw reads using kvarQ^67^.

### Phylogenetic reconstruction

The maximum likelihood phylogeny was inferred with RAxML v.8.2.8^68^ using an alignment containing only polymorphic sites and the branch lengths of the tree were rescaled using invariant sites (rescaled_branch_length = (branch_length * alignment_length) / (alignment_length +invariant_sites))^44,69^.

A position was considered polymorphic if at least one genome had a SNP at that position. Deletions and positions not called according to the minimum threshold of 7x were encoded as gaps. We excluded positions with more than 20% missing data, positions falling in PE-PGRS genes, phages, insertion sequences and in regions with at least 50 bp identity to other regions in the genome. We also excluded variable positions falling in drug resistance-related genes. The phylogeny was computed using the general time-reversible model of sequence evolution (-m GTRCAT-V options), 100 bootstrap inferences and *M. canettii* (SRR011186) was used as an outgroup to root the phylogeny.

### Whole genome *de novo* assembly, annotation and comparative genomics

Raw PacBio reads obtained from the Rwandan strain were assembled with Canu v1.6^70^, using default settings and an expected genome size of 4.4 Mbp, typical of MTBC strains. After discarding 60,272 reads below minimal quality parameters, 106,681 reads were used for the assembly. Based on the expected genome size, the average coverage depth was estimated at 186x using raw reads, and 39x and 38x using corrected and trimmed reads, respectively. The obtained unique contig of 4,387,285 bp was circularized with Circlator v1.5.5^71^ using default settings, resulting in an assembly of 4,379,493 bp. Additional sequence verification and correction was then performed by mapping Illumina reads obtained from the same strain, using pacbio-utils version 0.2^72^ (https://github.com/douglasgscofield/PacBio-utilities) and snippy version 4.4^73^ (https://github.com/tseemann/snippy). Alignments of the final assembly were performed against an ensemble of complete genome sequences available from 38 strains of tubercle bacilli. This set included 34 *M. tuberculosis* strains from lineages 1, 2, 3 and 4 (comprising H37Rv), *M. africanum* L6 GM041182, *M. bovis* AF2122/97, as well as the closest STB-A (CIPT 140010059) and most distant (STB-K) *M. canettii* strains (Supplementary Table 1). Comparative alignments and genome annotation were performed based on BLAST searches and analysis of gene synteny, using Artemis and Artemis comparison tool^74^, as well as a custom Multiple Annotation of Genomes and Differential Analysis (MAGDA) software previously used for annotation of *M. canettii* and Helicobacter pylori genomes^4,75^. Comparisons with orthologues from *M. canettii* STB-D, −E, −G, −H, −I, and −J in addition to STB-A and −K, and from *M. marinum* type strain M and *M. kansasii* genomes were additionally done using the Microscope platform v3.13.3^76^. When applicable, annotations were transferred from those of *M. tuberculosis* or *M. canettii* orthologs in the TubercuList/Mycobrowser database, using BLAST matches of > 90% protein sequence identity, an alignable region of >80% of the shortest protein length in pairwise comparisons and visual inspection of the gene synteny. Genome completeness was assessed using CheckM^77^ using the lineage-specific workflow and Mycobacterium as the genus. The PacBio assembled Rwandan strain was found to have 98.74% completeness and 0% contamination or strain heterogeneity, making it suitable for further analyses.

ACT comparison files were generated using MAUVE 2015-02-25 software to visualize the genome-wide distribution of SNP densities between the assembled L8 genome from Rwanda and *M. tuberculosis* H37Rv and *M. canettii* STB-A and STB-K genomes. Recombination between L8 and other MTBC lineages or *M. canettii* was assessed from a progressive MAUVE alignment of the PacBio assembled L8 genome and previously published closed genomes^73^ using ClonalFrameML^40^. To further assess the phylogenetic placement of the Lineage 8 strain, a core gene alignment was constructed using the completed genomes of the Rwandan L8 strain, representatives of the MTBC lineages 1-4 and 6, *M. canettii*, *M. bovis*, *M. marinum* (accession GCF_000419315.1) and M. kansassii (accession GCF_000157895.3). The GFF files of each genome were input to roary^78^ with an 80% identity cut-off, as has been done for previous genus-level mycobacterial core alignments^79^. A phylogenetic tree was constructed from the core gene superalignment using RAxML-NG v 0.9.0^80^ under the GTR+Gamma model of evolution with 20 starting trees. Bootstrapping was run until autoMRE converged with a value of 0.03 (50 replicates).

## Accession codes

The complete genome sequence of the L8 strain from Rwanda was deposited in the NCBI repository under project PRJNA598991 with SRR10828835 and SRR10828834 accession codes for Illumina- and PacBio-derived genome sequences, respectively.

## Acknowledgements

Calculations were partially performed at sciCORE (http://scicore.unibas.ch/) scientific computing core facility at University of Basel. This work was supported by EDCTP2 grant DRIA2014-326— DIAMA of the European Union, the Belgian General Directorate for Development Cooperation (PhD fellowship to JCSN), grant ANR-16-CE35-0009 from Agence Nationale de la Recherche, the Swiss National Science Foundation (grants 310030_188888, IZRJZ3_164171, IZLSZ3_170834 and CRSII5_177163), and the European Research Council (309540-EVODRTB). The views and opinions of authors expressed herein do not necessarily state or reflect those of EDCTP. P.S., C.G. and S.D. declare the following competing interests: P.S. was a consultant of Genoscreen; C.G. and S.D. were employees of the same company. The other authors declare no competing interests. The funders had no role in study design, data collection and analysis, decision to publish or preparation of the manuscript.

## Author contributions

S.G., P.S., C.M., B.C.d.J., L.R. and J.C.S.N. designed the study. P.S., J.C.S.N., C.L., M.M., C.M. and S.G. analyzed data and wrote the manuscript, with comments from all authors. A.J. and C.M. performed the assembly of sequences. M.M. annotated the L8 genome. C.L., F.M., D.B. and A.J. performed SNP analyses and phylogenetic reconstruction. M.M., A.R. and P.S. conducted comparative analyses of complete mycobacterial genomes, with support from C.L. and O.T. J.C.S.N., E.B.N., I.M.H., J.B.M., W.M., K.F. and M.D. performed and/or analyzed data from mycobacterial isolation, growth assays, phenotypic characterization and/or molecular tests. S.D., C.G., J.C.S.N., E.B.N., E.A. and M.K.K. conducted targeted deep sequencing analyses. S.D., C.G. and W.S. L. Majlessi, F.S., C. Locht and C. Leclerc conducted and/or analyzed immune assays. J.T., A. Criscuolo and S.B. conducted MLST, recombination and/or phylogenetic analyses. L.F. conducted histopathological analyses. V.K., M.O. and C.P. created bioinformatics tools and analyzed data. M.F. isolated STB strains. T.S. and T.P.S. conducted core genome and NeighborNet analyses.

## References

1. Gagneux, S. Ecology and evolution of Mycobacterium tuberculosis. Nature Reviews Microbiology 16, 202–213 (2018).

2. World Health Organization. Global Tuberculosis Report 2018. (2018).

3. Comas, I. et al. Out-of-Africa migration and Neolithic coexpansion of Mycobacterium tuberculosis with modern humans. Nature genetics 45, 1176–82 (2013).

4. Supply, P. et al. Genomic analysis of smooth tubercle bacilli provides insights into ancestry and pathoadaptation of Mycobacterium tuberculosis. Nature genetics 45, 172–9 (2013).

5. Bos, K. I. et al. Pre-Columbian mycobacterial genomes reveal seals as a source of New World human tuberculosis. Nature 514, 494–7 (2014).

6. Brites, D. et al. A New Phylogenetic Framework for the Animal-Adapted Mycobacterium tuberculosis Complex. Frontiers in Microbiology 9, 2820 (2018).

7. Rutaihwa, L. K. et al. Multiple Introductions of Mycobacterium tuberculosis Lineage 2– Beijing Into Africa Over Centuries. Frontiers in Ecology and Evolution 7, 112 (2019).

8. O’Neill, M. B. et al. Lineage specific histories of Mycobacterium tuberculosis dispersal in Africa and Eurasia. Molecular Ecology 28, 3241–3256 (2019).

9. Wirth, T. et al. Origin, spread and demography of the Mycobacterium tuberculosis complex. PLoS pathogens 4, e1000160 (2008).

10. Gagneux, S. et al. Variable host-pathogen compatibility in Mycobacterium tuberculosis. Proceedings of the National Academy of Sciences of the United States of America 103, 2869–73 (2006).

11. Gutierrez, M. C. et al. Ancient origin and gene mosaicism of the progenitor of Mycobacterium tuberculosis. PLoS pathogens 1, e5 (2005).

12. Blouin, Y. et al. Progenitor “Mycobacterium canettii” clone responsible for lymph node tuberculosis epidemic, Djibouti. Emerging infectious diseases 20, 21–8 (2014).

13. Coscolla, M. & Gagneux, S. Consequences of genomic diversity in Mycobacterium tuberculosis. Seminars in Immunology 26, 431–444 (2014).

14. Boritsch, E. C. et al. Key experimental evidence of chromosomal DNA transfer among selected tuberculosis-causing mycobacteria. Proceedings of the National Academy of Sciences of the United States of America 113, 9876–81 (2016).

15. Boritsch, E. C. et al. pks5-recombination-mediated surface remodelling in Mycobacterium tuberculosis emergence. Nature Microbiology 1, 15019 (2016).

16. Mortimer, T. D. & Pepperell, C. S. Genomic signatures of distributive conjugal transfer among mycobacteria. Genome biology and evolution 6, 2489–500 (2014).

17. Helb, D. et al. Rapid detection of Mycobacterium tuberculosis and rifampin resistance by use of on-demand, near-patient technology. Journal of Clinical Microbiology 48, 229–237 (2010).

18. Ng, K. C. S. et al. Automated algorithm for early identification of rifampicin-resistant tuberculosis transmission hotspots in Rwanda [abstract]. The International Journal of Tuberculosis and Lung Disease 22, 605 (2018).

19. Torrea, G. et al. Variable ability of rapid tests to detect Mycobacterium tuberculosis rpoB mutations conferring phenotypically occult rifampicin resistance. Scientific Reports 9, (2019).

20. Trébucq, A. et al. Treatment outcome with a short multidrug-resistant tuberculosis regimen in nine African countries. The International Journal of Tuberculosis and Lung Disease 22, 17–25 (2018).

21. Coll, F. et al. A robust SNP barcode for typing Mycobacterium tuberculosis complex strains. Nature Communications 5, (2014).

22. Couvin, D., David, A., Zozio, T. & Rastogi, N. Macro-geographical specificities of the prevailing tuberculosis epidemic as seen through SITVIT2, an updated version of the Mycobacterium tuberculosis genotyping database. Infection, Genetics and Evolution (2018). doi:10.1016/j.meegid.2018.12.030

23. Wanzala, S. I. et al. Retrospective Analysis of Archived Pyrazinamide Resistant Mycobacterium tuberculosis Complex Isolates from Uganda-Evidence of Interspecies Transmission. Microorganisms 7, (2019).

24. Stinear, T. P. et al. Insights from the complete genome sequence of Mycobacterium marinum on the evolution of Mycobacterium tuberculosis. Genome Research 18, 729–741 (2008).

25. Wang, J. et al. Insights on the emergence of Mycobacterium tuberculosis from the analysis of Mycobacterium kansasii. Genome biology and evolution 7, 856–70 (2015).

26. Ssengooba, W. et al. Whole genome sequencing to complement tuberculosis drug resistance surveys in Uganda. Infection, genetics and evolutionl⍰: journal of molecular epidemiology and evolutionary genetics in infectious diseases 40, 8–16 (2016).

27. Carpels, G. et al. Drug resistant tuberculosis in sub-Saharan Africa: an estimation of incidence and cost for the year 2000. Tubercle and lung diseasel⍰: the official journal of the International Union against Tuberculosis and Lung Disease 76, 480–6 (1995).

28. Umubyeyi, A. N. et al. Results of a national survey on drug resistance among pulmonary tuberculosis patients in Rwanda. The international journal of tuberculosis and lung diseaselZ: the official journal of the International Union against Tuberculosis and Lung Disease 11, 189–94 (2007).

29. World Health Organization & WHO. WHO | Global tuberculosis report 2016. WHO (2019).

30. Yang, T. et al. Pan-genomic study of Mycobacterium tuberculosis reflecting the primary/secondary genes, generality/individuality, and the interconversion through copy number variations. Frontiers in Microbiology 9, (2018).

31. Brosch, R. et al. A new evolutionary scenario for the Mycobacterium tuberculosis complex. Proceedings of the National Academy of Sciences of the United States of America 99, 3684–9 (2002).

32. Tsolaki, A. G. et al. Functional and evolutionary genomics of Mycobacterium tuberculosis: Insights from genomic deletions in 100 strains. Proceedings of the National Academy of Sciences of the United States of America 101, 4865–4870 (2004).

33. Brosch, R. et al. Genomic analysis reveals variation between Mycobacterium tuberculosis H37Rv and the attenuated M. tuberculosis H37Ra strain. Infection and Immunity 67, 5768–5774 (1999).

34. Hirsh, A. E., Tsolaki, A. G., DeRiemer, K., Feldman, M. W. & Small, P. M. Stable association between strains of Mycobacterium tuberculosis and their human host populations. Proceedings of the National Academy of Sciences of the United States of America 101, 4871–6 (2004).

35. Chiner-Oms et al. Genomic determinants of speciation and spread of the Mycobacterium tuberculosis complex. Science Advances 5, (2019).

36. Deshayes, C. et al. Detecting the molecular scars of evolution in the Mycobacterium tuberculosis complex by analyzing interrupted coding sequences. BMC evolutionary biology 8, 78 (2008).

37. Smith, N. H., Hewinson, R. G., Kremer, K., Brosch, R. & Gordon, S. V. Myths and misconceptions: the origin and evolution of Mycobacterium tuberculosis. Nature Reviews Microbiology 7, 537–544 (2009).

38. Etienne, G. et al. Identification of the polyketide synthase involved in the biosynthesis of the surface-exposed lipooligosaccharides in mycobacteria. Journal of Bacteriology 191, 2613–2621 (2009).

39. Boritsch, E. C. & Brosch, R. Evolution of Mycobacterium tuberculosis: New Insights into Pathogenicity and Drug Resistance. in Tuberculosis and the Tubercle Bacillus, Second Edition 4, 495–515 (American Society of Microbiology, 2016).

40. Didelot, X. & Wilson, D. J. ClonalFrameML: efficient inference of recombination in whole bacterial genomes. PLoS computational biology 11, e1004041 (2015).

41. Hershberg, R. et al. High Functional Diversity in Mycobacterium tuberculosis Driven by Genetic Drift and Human Demography. PLoS Biology 6, e311 (2008).

42. Supply, P. et al. Linkage disequilibrium between minisatellite loci supports clonal evolution of Mycobacterium tuberculosis in a high tuberculosis incidence area. Molecular Microbiology 47, 529–538 (2003).

43. Mostowy, S., Cousins, D., Brinkman, J., Aranaz, A. & Behr, M. A. Genomic Deletions Suggest a Phylogeny for the *Mycobacterium tuberculosis* Complex. The Journal of Infectious Diseases 186, 74–80 (2002).

44. Menardo, F., Duchêne, S., Brites, D. & Gagneux, S. The molecular clock of Mycobacterium tuberculosis. PLoS pathogens 15, e1008067 (2019).

45. Niobe-Eyangoh, S. N. et al. Genetic biodiversity of Mycobacterium tuberculosis complex strains from patients with pulmonary tuberculosis in Cameroon. Journal of Clinical Microbiology 41, 2547–2553 (2003).

46. Godreuil, S. et al. First molecular epidemiology study of Mycobacterium tuberculosis in Burkina Faso. Journal of Clinical Microbiology 45, 921–927 (2007).

47. Groenheit, R. et al. The Guinea-Bissau Family of Mycobacterium tuberculosis Complex Revisited. PLoS ONE 6, e18601 (2011).

48. Comas, I. et al. Population Genomics of Mycobacterium tuberculosis in Ethiopia Contradicts the Virgin Soil Hypothesis for Human Tuberculosis in Sub-Saharan Africa. Current Biology 25, 3260–3266 (2015).

49. Ates, L. S. et al. Mutations in ppe38 block PE_PGRS secretion and increase virulence of Mycobacterium tuberculosis. Nature Microbiology 3, 181–188 (2018).

50. Boritsch, E. C. et al. A glimpse into the past and predictions for the future: the molecular evolution of the tuberculosis agent. Molecular Microbiology 93, 835–852 (2014).

51. Martens, J. H., Barg, H., Warren, M. & Jahn, D. Microbial production of vitamin B12. Applied Microbiology and Biotechnology 58, 275–285 (2002).

52. Gopinath, K. et al. A vitamin B_12_ transporter in *Mycobacterium tuberculosis*. Open Biology 3, 120175 (2013).

53. Minias, A., Minias, P., Czubat, B. & Dziadek, J. Purifying Selective Pressure Suggests the Functionality of a Vitamin B12 Biosynthesis Pathway in a Global Population of Mycobacterium tuberculosis. Genome biology and evolution 10, 2326–2337 (2018).

54. Bottai, D. et al. TbD1 deletion as a driver of the evolutionary success of modern epidemic Mycobacterium tuberculosis lineages. Nature Communications 11, 1–14 (2020).

55. Current BiologyLEÃwO, S. C. et al. Practical handbook for the phenotypic and genotypic identification of mycobacteria. (2004).

56. Borrell, S. et al. Reference set of Mycobacterium tuberculosis clinical strains: A tool for research and product development. PLOS ONE 14, e0214088 (2019).

57. Kent, P. & Kubica, G. Public Health Mycobacteriology: A Guide for the Level III Laboratory US Department of Health and Human Services, Centres for Disease Control. (1985).

58. Makhado, N. A. et al. Outbreak of multidrug-resistant tuberculosis in South Africa undetected by WHO-endorsed commercial tests: an observational study. The Lancet Infectious Diseases 18, 1350–1359 (2018).

59. van der Zanden, A. G. M. et al. Improvement of differentiation and interpretability of spoligotyping for Mycobacterium tuberculosis complex isolates by introduction of new spacer oligonucleotides. Journal of Clinical Microbiology 40, 4628–39 (2002).

60. Menardo, F. et al. Treemmer: a tool to reduce large phylogenetic datasets with minimal loss of diversity. BMC Bioinformatics 19, 164 (2018).

61. Bolger, A. M., Lohse, M. & Usadel, B. Trimmomatic: a flexible trimmer for Illumina sequence data. Bioinformatics (Oxford, England) 30, 2114–20 (2014).

62. Comas, I. et al. Human T cell epitopes of Mycobacterium tuberculosis are evolutionarily hyperconserved. Nature genetics 42, 498–503 (2010).

63. Li, H. & Durbin, R. Fast and accurate short read alignment with Burrows-Wheeler transform. Bioinformatics (Oxford, England) 25, 1754–60 (2009).

64. McKenna, A. et al. The Genome Analysis Toolkit: a MapReduce framework for analyzing next-generation DNA sequencing data. Genome Research 20, 1297–303 (2010).

65. Li, H. A statistical framework for SNP calling, mutation discovery, association mapping and population genetical parameter estimation from sequencing data. Bioinformatics (Oxford, England) 27, 2987–93 (2011).

66. Koboldt, D. C. et al. VarScan 2: somatic mutation and copy number alteration discovery in cancer by exome sequencing. Genome Research 22, 568–76 (2012).

67. Steiner, A., Stucki, D., Coscolla, M., Borrell, S. & Gagneux, S. KvarQ: targeted and direct variant calling from fastq reads of bacterial genomes. BMC Genomics 15, 881 (2014).

68. Stamatakis, A. RAxML version 8: a tool for phylogenetic analysis and post-analysis of large phylogenies. Bioinformatics 30, 1312–1313 (2014).

69. Duchene, S. et al. Inferring demographic parameters in bacterial genomic data using Bayesian and hybrid phylogenetic methods. BMC evolutionary biology 18, 95 (2018).

70. Koren, S. et al. Canu: scalable and accurate long-read assembly via adaptive k-mer weighting and repeat separation. Genome Research 27, 722–736 (2017).

71. Hunt, M. et al. Circlator: automated circularization of genome assemblies using long sequencing reads. Genome Biology 16, 294 (2015).

72. Scofield, D. G. GitHub - douglasgscofield/PacBio-utilities: Collection of utilities for working with PacBio-based assemblies. Available at: https://github.com/douglasgscofield/PacBio-utilities. (Accessed: 20th December 2019)

73. Seemann, T. GitHub - tseemann-snippy Rapid bacterial SNP calling and core genome alignments. Available at: https://github.com/tseemann/snippy. (Accessed: 20th December 2019)

74. Carver, T. et al. Artemis and ACT: viewing, annotating and comparing sequences stored in a relational database. Bioinformatics (Oxford, England) 24, 2672–6 (2008).

75. Li, H. et al. East-Asian Helicobacter pylori strains synthesize heptan-deficient lipopolysaccharide. PLoS Genetics 15, (2019).

76. Médigue, C. et al. MicroScope-an integrated resource for community expertise of gene functions and comparative analysis of microbial genomic and metabolic data. Briefings in bioinformatics 20, 1071–1084 (2019).

77. Parks, D. H., Imelfort, M., Skennerton, C. T., Hugenholtz, P. & Tyson, G. W. CheckM: Assessing the quality of microbial genomes recovered from isolates, single cells, and metagenomes. Genome Research 25, 1043–1055 (2015).

78. Page, A. J. et al. Roary: Rapid large-scale prokaryote pan genome analysis. Bioinformatics 31, 3691–3693 (2015).

79. Fedrizzi, T. et al. Genomic characterization of Nontuberculous Mycobacteria. Scientific Reports 7, (2017).

80. Kozlov, A. M., Darriba, D., Flouri, T., Morel, B. & Stamatakis, A. RAxML-NG: a fast, scalable and user-friendly tool for maximum likelihood phylogenetic inference. Bioinformatics (Oxford, England) 35, 4453–4455 (2019).

