## Supplementary Methods for "A sister lineage of the *Mycobacterium tuberculosis complex* discovered in the African Great Lakes region"

**Supplementary method 1: DNA extraction for targeted and Illumina sequencing**

A loopful colonies from LJ were suspended in Tris-EDTA buffer, and heat inactivated at 95°C for 20min. After cooling at room temperature, the suspension was centrifuged at 20,000g for 30 minutes followed by discarding supernatant and 250 µl of 10 mM Tris-HCl pH 7.8 were added and briefly vortexed. Mixture was incubated at 95°C for 15 minutes, then spun down briefly followed by transferring entire volume in a new microcentrifuge tube containing 0.5 g of zirconium beads (Sigma-Aldrich, St. Louis, USA). For destruction of the solid mycobacterial cell wall, the mixture was vortexed at high speed for at least 30 seconds followed by briefly spinning down and incubation at -20°C for at least 30 minutes. After thawing at room temperature, the mixture was briefly spun down and 200µl of supernatant was transferred to a new microcentrifuge tube. For gDNA concentration, 1µl of glycogen solution (Sigma-Aldrich, St. Louis, USA) was added and mixed followed by adding 0.1 volume of 3M sodium acetate at pH 5.2 (Thermo Fisher Scientific, Waltham, MA USA) and mixed, then 3 volume of 100% pre-cooled ethanol were added and vigorously vortexed for 10 seconds. The mixture was incubated at -20°C for 10 minutes. After thawing at room temperature, the mixture was centrifuged at 15000g for 20 minutes followed by discarding supernatant, then 600µl of freshly prepared pre-cooled 70% ethanol was added followed by centrifugation at 15000g for 5 minutes. Supernatant was discarded and the tube was air dried. gDNA was resuspended in 20µl of sterile molecular grade water. The yield was measured by Qubit dsDNA BR Assay Kit (Life Technologies, Carlsbad, USA).

**Supplementary method 2: DNA extraction for PacBio sequencing**

Colonies (70,3 ± 1,0 mg) from a one month old LJ were transferred into a 50ml falcon tube containing 3,5 ml of the Qiagen buffer B1 and 70µL of 10 mg/ml RNAse (Life Technologies, Carlsbad, USA) solution followed by thoroughly vortex and heat inactivation of bacilli at 80°C for 1 hour. After cooling at room temperature, 100 µL of 100 mg/ml lysozyme (Sigma-Aldrich, St. Louis, USA) was added, followed by inverting at least 20 times, then placed in horizontal shaker at 37°C (0 RPM) for 60 min. After, 1.2 ml of 2.5 mg/ml of Proteinase K (MP Biomedicals, Santa Ana, USA) was added followed by inverting tube at least 20 times, then placed in horizontal shaker at 37°C (0 RPM) for 60 min. For protein denaturation (nucleases and DNA-binding proteins), 1.2 ml of buffer B2 was added to the mixture and placed overnight in horizontal shaker at 50°C (30 RPM) alongside with Qiagen buffer QF. On the next day, the mixture was swirled gently by hand for 1 min and centrifuged at 3700g, 4°C for 10min after adding 4 ml of buffer QBT to a 100/G tip to equilibrate and allowed it to flow through, then the supernatant from centrifugated mixture tube was loaded by tipping onto the column and allowing the supernatant to flow through, and the cap of the mixture tube was placed on the column followed by adding 7.5 ml of buffer QC to the tip holder. Column was placed over a new clean pre-labelled 50ml falcon tube, and the DNA was eluted in 5ml pre-warmed QF buffer. For DNA precipitation, 3.5 ml of isopropanol were added and mixed by inverting 20 times followed by centrifugation at 4500g, 4°C for 20 min, then the supernatant was gently discarded. gDNA was then suspended in 1.5 ml cold (4°C) 70% ethanol and pellet washed by gently swirling the tube by hand. The entire volume was transferred to 1.5 ml tube and centrifuged at 10.000g, 4°C for 15 min followed by discarding all supernatant and drying the pellet in the Speedvac for 5 min at the medium drying setting. Finally, 200µl of 10mM Tris-Cl, pH 8,0 buffer was added and mixed gently by tapping and refrigeration overnight. On the next day, gDNA pellet was resuspended in the horizontal shaker at 65°C (250 RPM) for 15 min. The purity and integrity of the gDNA were examined through 0.5% agarose gel electrophoresis and yield was measured using the NanoDrop 1000 Spectrophotometer (Thermo Fisher Scientific, Waltham, MA USA) and Qubit dsDNA BR Assay Kit (Life Technologies, Carlsbad, USA) according to manufacturer’s instructions.
