## Supplementary Table 1 for "A sister lineage of the *Mycobacterium tuberculosis complex* discovered in the African Great Lakes region"

Supplementary 1. Information on the 38 complete genome sequences of MTBC and *M. canettii* strains used in this study (adapted from Yang; T., Zhong, J., Zhang, J., Cuidan, L., Yu, X., Xiao, J. et al. (2018). Pan-genomic study of Mycobacterium tuberculosis reflecting the primary/secondary genes, generality/individuality, and the interconversion through copy number variations. *Front. Microbiol.* 9. doi: 10.3389/fmicb.2018.01886).

| **Strains** | **Accession No.** | **ATCC No./Lineage** | **Genome size (Mb)** | **GC content (%)** | **Coding gene No.** | **% Coding region** | **Source [reference]** |
| --- | --- | --- | --- | --- | --- | --- | --- |
| *M. tuberculosis* H37Rv | NC_000962 | L4 | 4.41 | 65.6 | 4336 | 90.42 | (Cole et al., 1998) |
| *M. tuberculosis* F1 | CP010329 | 27294/L4 | 4.43 | 65.61 | 4400 | 90.26 | Beijing Institute of Genomics (Zhu et al., 2016) |
| *M. tuberculosis* F28 | CP010330 | 25177/L4 | 4.42 | 65.6 | 4366 | 90.26 | Beijing Institute of Genomics (Zhu et al., 2016) |
| *M. tuberculosis* H37Ra | NC_009525 | 25177/L4 | 4.42 | 65.6 | 4348 | 90.4 | Fudan University (Zheng et al., 2008) |
| *M. tuberculosis* Erdman | NC_020559 | 35801/L4 | 4.39 | 65.6 | 4373 | 90.34 | Tohru Akiyama National Center for Global Health and Medicine (Miyoshi-Akiyama et al., 2012) |
| *M. tuberculosis* 22103 | CP010339 | L4 | 4.4 | 65.61 | 4345 | 90.23 | Beijing Institute of Genomics (Zhu et al., 2016) |
| *M. tuberculosis* 22115 | CP010337 | L4 | 4.4 | 65.57 | 4356 | 90.23 | Beijing Institute of Genomics (Zhu et al., 2016) |
| *M. tuberculosis* 37004 | CP010338 | L4 | 4.42 | 65.6 | 4375 | 90.28 | Beijing Institute of Genomics (Zhu et al., 2016) |
| *M. tuberculosis* KZN 4207 | NC_016768 | L4 | 4.4 | 65.6 | 4324 | 90.42 | Broad Institute of MIT and Harvard (Ioerger et al., 2009) |
| *M. tuberculosis* KZN 605 | NC_018078 | L4 | 4.4 | 65.6 | 4326 | 90.31 | Broad Institute of MIT and Harvard (Ioerger et al., 2009) |
| *M. tuberculosis* KZN 1435 | NC_012943 | L4 | 4.4 | 65.6 | 4333 | 90.36 | Broad Institute of MIT and Harvard (Ioerger et al., 2009) |
| *M. tuberculosis* Haarlem | NC_022350 | L4 | 4.41 | 65.6 | 4322 | 90.39 | Broad Institute of MIT and Harvard (Mardassi et al., 2005) |
| *M. tuberculosis* F11 | NC_009565 | L4 | 4.42 | 65.6 | 4352 | 90.41 | Broad Institute of MIT and Harvard (Victor et al., 2004) |
| *M. tuberculosis* CDC1551 | NC_002755 | L4 | 4.4 | 65.6 | 4351 | 90.33 | The Institute for Genomic Research (Fleischmann et al., 2002) |
| *M. tuberculosis* 7199-99 | NC_020089 | L4 | 4.42 | 65.6 | 4344 | 90.47 | Bielefeld University (Roetzer et al., 2013) |
| *M. tuberculosis* CTRI-2 | NC_017524 | L4 | 4.4 | 65.6 | 4331 | 90.39 | Research Institute of Physical-Chemical Medicine (Ilina et al., 2013) |
| *M. tuberculosis* Kurono | NZ_AP014573 | 35812/L4 | 4.42 | 65.6 | 4342 | 90.4 | Department of Infectious Disease, Japan (Miyoshi-Akiyama et al., 2015) |
| *M. tuberculosis* 26105 | CP010340 | L3 | 4.43 | 65.63 | 4393 | 90.29 | Beijing Institute of Genomics (Zhu et al., 2016) |
| *M. tuberculosis* NITR203 | NC_021054 | L2 | 4.41 | 65.6 | 4442 | 90.32 | National Institute of Research in Tuberculosis, India (Narayanan and Deshpande., 2013) |
| *M. tuberculosis* HKBS1 | CP002871 | L2 | 4.41 | 65.6 | 4343 | 90.44 | Chinese University of Hong Kong |
| *M. tuberculosis* CCDC5079 | NC_021251 | L2 | 4.41 | 65.6 | 4354 | 90.39 | Chinese Center for Disease Control and Prevention (Tang et al., 2013) |
| *M. tuberculosis* 49-02 | HG813240 | L2 | 4.42 | 65.6 | 4350 | 90.48 | Justus-Liebig-University Giessen |
| *M. tuberculosis* 96075 | CP009426 | L2 | 4.4 | 65.6 | 4327 | 90.44 | University of Hawaii (Wan et al., 2014) |
| *M. tuberculosis* BT1 | CP002883 | L2 | 4.4 | 65.6 | 4337 | 90.44 | Chinese University of Hong Kong |
| *M. tuberculosis* BT2 | CP002882 | L2 | 4.4 | 65.6 | 4343 | 90.34 | Chinese University of Hong Kong |
| *M. tuberculosis* CCDC5180 | CP002885 | L2 | 4.41 | 65.6 | 4353 | 90.4 | Chinese Center for Disease Control and Prevention |
| *M. tuberculosis* 323 | CP010873 | L2 | 4.41 | 65.6 | 4403 | 90.41 | (Rodriguez et al., 2015) |
| *M. tuberculosis* ZMC13-88 | CP009101 | L2 | 4.41 | 65.6 | 4364 | 90.52 | Affiliated Hospital of Zunyi Medical College (Chen et al., 2014) |
| *M. tuberculosis* ZMC13-264 | CP009100 | L2 | 4.41 | 65.6 | 4365 | 90.46 | Affiliated Hospital of Zunyi Medical College (Chen et al., 2014) |
| *M. tuberculosis* KIT87190 | CP007809 | L2 | 4.41 | 65.6 | 4341 | 90.35 | Korean Institute of Tuberculosis (Park et al., 2014) |
| *M. tuberculosis* K | CP007803 | L2 | 4.39 | 65.59 | 4334 | 90.45 | Yonsei University College of Medicine (Ryoo et al., 2007) |
| *M. tuberculosis* 96121 | CP009427 | L1 | 4.41 | 65.6 | 4375 | 90.37 | University of Hawaii (Wan et al., 2014) |
| *M. tuberculosis* EAI5 | NC_021740 | L1 | 4.39 | 65.6 | 4322 | 90.5 | University of Mumbai (Al Rashdi et al., 2014) |
| *M. tuberculosis* NITR206 | NC_021194 | L1 | 4.39 | 65.6 | 4417 | 90.32 | National Instute of Research in Tuberculosis, India (Narayanan and Deshpande., 2013) |
| *M. bovis* AF/2227 |  | Animal |  |  |  |  | (Garnier et al., 2003) |
| *M. africanum* GM041182 |  | L6 |  |  |  |  | (Bentley et al., 2012) |
| *M. canettii* STB-A |  |  |  |  |  |  | (Supply et al., 2013) |
| *M. canettii* STB-K |  |  |  |  |  |  | (Supply et al., 2013) |

**References**

Al Rashdi, A. S., Jadhav, B. L., Deshpande, T., and Deshpande, U. (2014). Whole-genome sequencing and annotation of a clinical isolate of *Mycobacterium tuberculosis* from mumbai, india. *Genome. Announc*. 2. doi: 10.1128/genomeA.00154-14.

Bentley SD, Comas I, Bryant JM, Walker D, Smith NH, Harris SR, et al. (2012). The genome of Mycobacterium africanum West African 2 reveals a lineage-specific locus and genome erosion common to the M. tuberculosis complex. *PLoS Negl. Trop. Dis.* 6(2):e1552. doi: 10.1371.

Chen, L., Zhang, D. T., Zhang, J., Su, Y. A., and Zhang, H. (2014). Whole-genome sequences of two clinical isolates of extensively drug-resistant *Mycobacterium tuberculosis* from Zunyi, China. *Genome. Announc*. 2. doi: 10.1128/genomeA.00910-14.

Cole, S. T., Brosch, R., Parkhill, J., Garnier, T., Churcher, C., Harris, D., et al. (1998). Deciphering the biology of *Mycobacterium tuberculosis* from the complete genome sequence. *Nature*. 393, 537-544.

Fleischmann, R. D., Alland, D., Eisen, J. A., Carpenter, L., White, O., Peterson, J., et al. (2002). Whole-genome comparison of *Mycobacterium tuberculosis* clinical and laboratory strains. *J. Bacteriol.* 184, 5479-5490. doi: 10.1128/Jb.184.19.5479-5490.2002.

Garnier, T., Eiglmeier, K., Camus, J.C., Medina, N., Mansoor, H., Pryor, M., Duthoy, S., Grondin, S., Lacroix, C., Monsempe, C., Simon, S., Harris, B., Atkin, R., Doggett, J., Mayes, R., Keating, L., Wheeler, P.R., Parkhill, J., Barrell, B.G., Cole S.T., Gordon S.V., Hewinson, R.G. (2003). The complete genome sequence of Mycobacterium bovis. *Proc. Natl Acad. Sci. U S A*. 100, 7877-7882.

Ilina, E. N., Shitikov, E. A., Ikryannikova, L. N., Alekseev, D. G., Kamashev, D. E., Malakhova, M. V., et al. (2013). Comparative genomic analysis of *Mycobacterium tuberculosis* drug resistant strains from Russia. *PLoS. One*. 8. doi: ARTN e56577

10.1371/journal.pone.0056577.

Ioerger, T. R., Koo, S., No, E. G., Chen, X., Larsen, M. H., Jacobs, W. R., Jr., et al. (2009). Genome analysis of multi- and extensively-drug-resistant tuberculosis from KwaZulu-Natal, South Africa. *PLoS. One*. 4, e7778. doi: 10.1371/journal.pone.0007778.

Mardassi, H., Namouchi, A., Haltiti, R., Zarrouk, M., Mhenni, B., Karboul, A., et al. (2005). Tuberculosis due to resistant Haarlem strain, Tunisia. *Emerg. Infect. Dis*. 11, 957-961.

Miyoshi-Akiyama, T., Matsumura, K., Iwai, H., Funatogawa, K., and Kirikae, T. (2012). Complete annotated genome sequence of *Mycobacterium tuberculosis* Erdman. *J. Bacteriol*. 194, 2770. doi: 10.1128/JB.00353-12.

Miyoshi-Akiyama, T., Satou, K., Kato, M., Shiroma, A., Matsumura, K., Tamotsu, H., et al. (2015). Complete annotated genome sequence of *Mycobacterium tuberculosis* (Zopf) Lehmann and neumann (ATCC35812) (Kurono). *Tuberculosis*. 95, 37-39. doi: 10.1016/j.tube.2014.10.007.

Narayanan, S., and Deshpande, U. (2013). Whole-genome sequences of four clinical isolates of mycobacterium tuberculosis from tamil nadu, south india. *Genome. Announc*. 1. doi: 10.1128/genomeA.00186-13.

Park, Y. K., Kang, H., Yoo, H., Lee, S. H., Roh, H., Kim, H. J., et al. (2014). Whole-genome sequence of *Mycobacterium tuberculosis* Korean strain KIT87190. *Genome. Announc*. 2. doi: 10.1128/genomeA.01103-14.

Rodriguez, J. G., Pino, C., Tauch, A., and Murcia, M. I. (2015). Complete genome sequence of the clinical Beijing-like strain *Mycobacterium tuberculosis* 323 using the PacBio real-time sequencing platform. *Genome. Announc*. 3. doi: 10.1128/genomeA.00371-15.

Roetzer, A., Diel, R., Kohl, T. A., Ruckert, C., Nubel, U., Blom, J., et al. (2013). Whole genome sequencing versus traditional genotyping for investigation of a *Mycobacterium tuberculosis* outbreak: A longitudinal molecular epidemiological study. *PLoS. Med*. 10. doi: ARTN e100138710.1371/journal.pmed.1001387.

Ryoo, S. W., Park, Y. K., Park, S. N., Shim, Y. S., Liew, H., Kang, S., et al. (2007). Comparative proteomic analysis of virulent Korean *Mycobacterium tuberculosis* K-strain with other mycobacteria strain following infection of U-937 macrophage. *J. Microbio.* 45, 268-271.

Supply, P., Marceau, M., Mangenot, S., Roche, D., Rouanet, C., Khanna, V., et al. (2013). Genomic analysis of smooth tubercle bacilli provides insights into ancestry and pathoadaptation of Mycobacterium tuberculosis. *Nat. Genet.* 45, 172-179.

Tang, B., Wang, Q., Yang, M., Xie, F., Zhu, Y., Zhuo, Y., et al. (2013). ContigScape: A Cytoscape plugin facilitating microbial genome gap closing. *BMC. Genomics*. 14, 289. doi: 10.1186/1471-2164-14-289.

Victor, T. C., de Haas, P. E. W., Jordaan, A. M., van der Spuy, G. D., Richardson, M., van Soolingen, D., et al. (2004). Molecular characteristics and global spread of *Mycobacterium tuberculosis* with a western cape F11 genotype. *J. Clin. Microbiol*. 42, 769-772. doi: 10.1128/Jcm.42.2.769-772.2004.

Wan, X., Qian, L., Hou, S., Drees, K. P., Foster, J. T., and Douglas, J. T. (2014). Complete genome sequences of Beijing and Manila family strains of *Mycobacterium tuberculosis*. *Genome. Announc*. 2. doi: 10.1128/genomeA.01135-14.

Zheng, H., Lu, L., Wang, B., Pu, S., Zhang, X., Zhu, G., et al. (2008). Genetic basis of virulence attenuation revealed by comparative genomic analysis of *Mycobacterium tuberculosis* strain H37Ra versus H37Rv. *PLoS. One.* 3, e2375. doi: 10.1371/journal.pone.0002375.

Zhu, L. X., Zhong, J., Jia, X. M., Liu, G., Kang, Y., Dong, M. X., et al. (2016). Precision methylome characterization of *Mycobacterium tuberculosis* complex (MTBC) using PacBio single-molecule real-time (SMRT) technology. *Nucleic. Acids. Res*. 44, 730-743. doi: 10.1093/nar/gkv1498.
