## Supplementary Figure 1 for "A sister lineage of the *Mycobacterium tuberculosis complex* discovered in the African Great Lakes region"

**Supplementary figure 1: Core genome-based cladogram of the MTBC and outgroups.** A phylogeny was constructed from the core alignment of L8, MTBC lineage representatives and *M. canettii* with *M. marinum* and *M. kansasii* as outgroups. Node labels represent bootstrap support for that split. Phylogeny is shown as a cladogram (branch lengths have no meaning) to allow clearer visualisation of the topology.

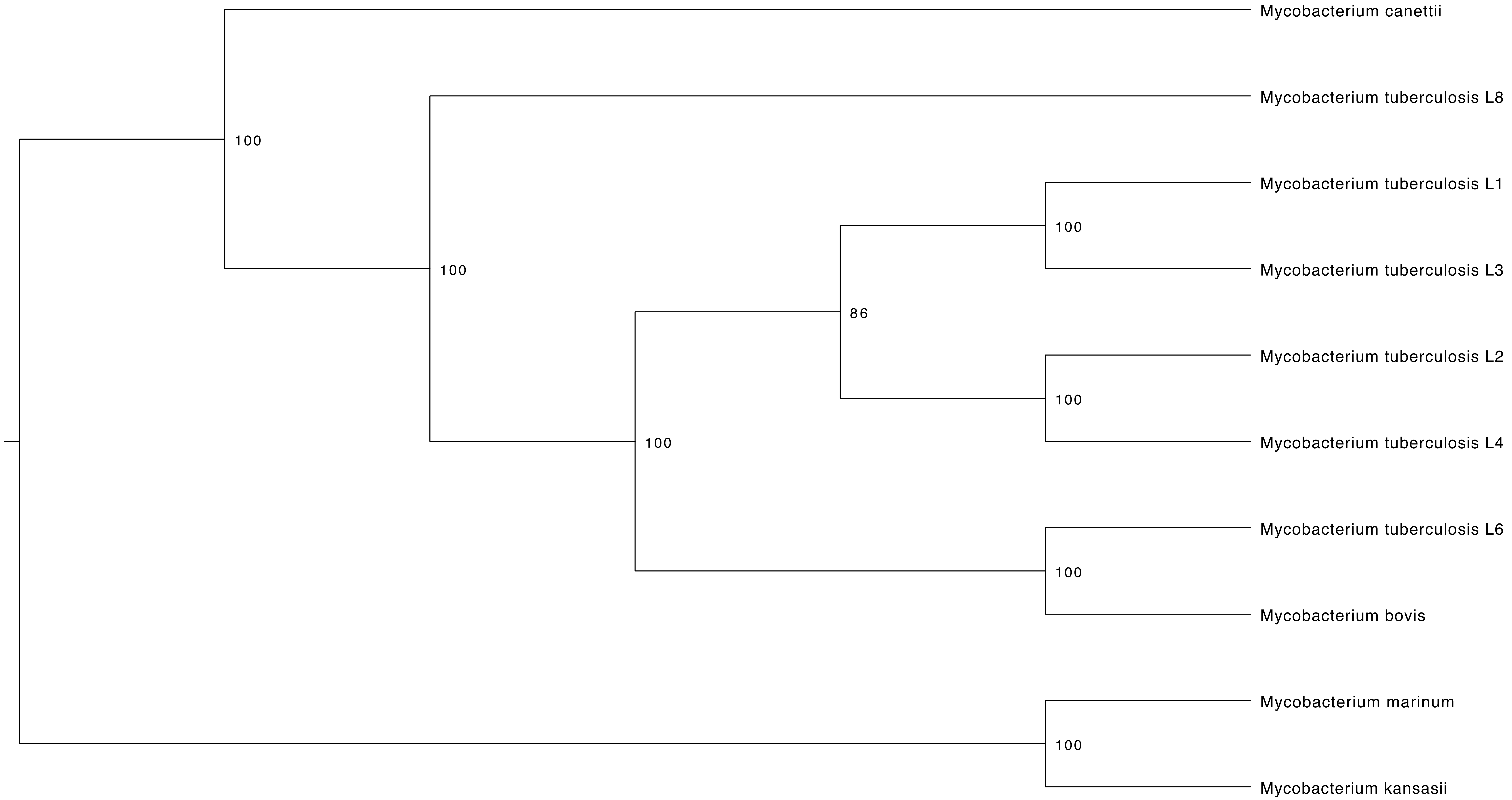
