## Supplementary Figure 2 for "A sister lineage of the *Mycobacterium tuberculosis complex* discovered in the African Great Lakes region"

**Supplementary figure 2: ClonalFrameML recombination analysis of the MTBC and *M. canettii*.** White bars indicate reconstructed substitutions. Dark blue dots indicate areas of recombination.

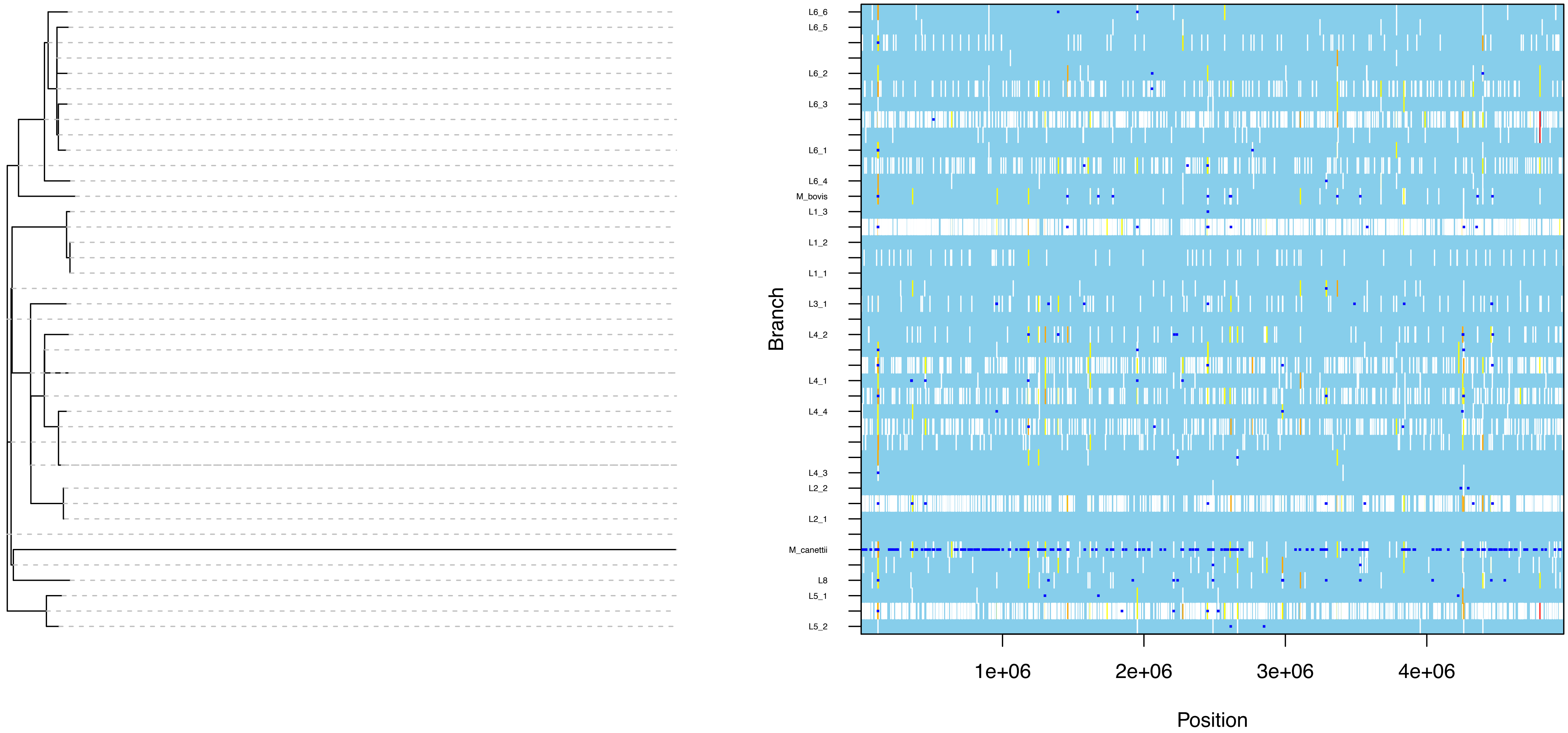
